# Webina: An Open-Source Library and Web App that Runs AutoDock Vina Entirely in the Web Browser

**DOI:** 10.1101/2019.12.18.881789

**Authors:** Yuri Kochnev, Erich Hellemann, Kevin C. Cassidy, Jacob D. Durrant

## Abstract

Molecular docking is a computational technique for predicting how a small molecule might bind a macromolecular target. Among docking programs, AutoDock Vina is particularly popular. Like many docking programs, Vina requires users to download/install an executable file and to run that file from a Unix- or DOS-like command-line interface. Choosing proper configuration parameters and analyzing Vina output is also sometimes challenging. These issues are particularly problematic for students and novice researchers. We have created Webina, a new version of Vina, to address these challenges. Webina is a JavaScript/WebAssembly library that runs AutoDock Vina entirely in a web browser. To use Webina, users need only visit a Webina-enabled webpage. The docking calculations take place on the user’s own computer rather than a remote server. To encourage use, we have incorporated the Webina library into our own Webina web app. The app includes a convenient interface so users can easily setup their docking runs and analyze the results. Webina will be a useful open-source tool for the research and educational communities. A working version of the app can be accessed free of charge from http://durrantlab.com/webina.

## 2 Introduction

Molecular docking is a popular computer-aided drug discovery (CADD) technique for predicting non-covalent small-molecule/macromolecular binding. By accelerating lead identification, docking aims to streamline the early-stage drug-discovery process. A docking program first predicts the 3D geometries (“poses”) with which virtual-library compounds might bind a given macromolecular target. Second, a scoring function evaluates the predicted geometries to estimate binding affinities [1]. The top-scoring compounds are then recommended for experimental testing. The resulting hit rates are far from 100%, but they tend to be better than those obtained from more costly high-throughput experimental screens [2].

Among docking programs, AutoDock Vina is particularly popular [1]. Vina is an open-source program written in C++ that runs on all major desktop operating systems. Its strengths include speed and relative ease of use. But like many CADD programs, Vina has some notable shortcomings. Users must download and install the program to use it on their own machines. Choosing proper configuration parameters and analyzing Vina output is also sometimes challenging. And absent third-party graphical user interface (GUI) wrappers [3–7], Vina is only accessible from a Unix- or DOS-like command-line interface. These limitations are particularly impactful in educational settings, where expecting students to download, install, and use a command-line program is often impractical.

To address these challenges, we have created Webina, a WebAssembly (Wasm) library that runs AutoDock Vina entirely in a web browser. Wasm is an emerging web technology that allows developers to run compiled code without requiring the installation of any third-party plugins or programs. High-level languages such as C, C++, and Rust can be compiled to Wasm just as they can be compiled to operating-system-specific binary executable files. But Wasm-compiled programs run on a browser-based virtual machine rather than on a desktop operating system. Wasm can thus transform a stand-alone computer program into a browser-compatible library that can be accessed via standard web applications. And because Wasm code runs CPU- and/or memory-intensive operations on the user’s own computer, those who create such web applications do not need to maintain the computer infrastructure typically required to run complex calculations on remote servers.

To create the Webina library, we compiled the AutoDock Vina 1.1.2 codebase to Wasm using Emscripten 1.38.48, a toolchain specifically designed to port C/C++ applications to the web (https://emscripten.org). Our benchmarks show that our optimized Webina Wasm code runs only ~10% slower than the original Vina binary.

To facilitate use, we also created a web app with a convenient GUI interface so users can easily configure and run Webina on their own receptor/ligand systems of interest. The same app also allows users to visualize Webina-docked poses in their browsers.

Webina will be a useful tool for the CADD research and educational communities. The web-app source code and compiled Webina library are available at http://durrantlab.com/webina-download. We release both under the terms of the open-source Apache License, Version 2.0. A working version of the app can be accessed free of charge from http://durrantlab.com/webina.

## 3 Materials and Methods

### 3.1 Webina Library: Compiling Vina to WebAssembly

We compiled Vina to Wasm without modifying the original Vina 1.1.2 source code. Fortunately, Vina has only one dependency: the popular Boost libraries for C++. Many of these libraries are header-only, so including them in Emscripten-compiled projects requires no preliminary compilation. But Vina directly calls Boost libraries that are not header-only. These had to be first recompiled as static libraries for Wasm before compiling the main project.

The age of the Vina source code presented a major challenge. Vina has not been modified since 2011 and so is compatible only with older versions of Boost. The latest version of Boost is 1.71, but we used the *system, filesystem, serialization, program_options and thread/pthread* libraries from Boost 1.41. Aside from overcoming small bugs mostly related to C++11 incompatibilities, the key to successful compilation was to provide Emscripten with the required Boost include files. The resulting binaries had to be linked to *em++* during the compilation process by modifying the Vina Makefile. Further details can be found in the README.md file included with the Webina download.

We have tested the Webina library on the browser/operating-system combinations shown in Table 1. Webina uses the SharedArrayBuffer JavaScript object to allow multiple processes/threads to exchange data directly. Most browsers disabled this object in 2018 due to concerns over the Spectre and Meltdown exploits. But it is currently available on Chromium-based browsers such as Google Chrome, and additional browsers (e.g., Firefox, Safari) are likely to reenable SharedArrayBuffer soon.

**Table 1:** Browser compatibility. We have tested Webina on Chromium-based browsers running on all major desktop operating systems.

| Browser | Operating System |
| --- | --- |
| Chrome 79.0.3945.36 | macOS 10.14.5 |
| Chrome 78.0.3904.97 | Windows 10 Home 1803 |
| Chromium 78.0.3904.70 | Ubuntu 18.04.3 LTS |

### 3.2 Webina Web App

#### 3.2.1 Typescript

To facilitate use, we have incorporated the Webina library into a convenient web app. The app is written in the open-source Microsoft TypeScript programming language. TypeScript is similar to JavaScript, but it includes additional features (e.g., optional variable typing, standard classes and interfaces, etc.) that are particularly useful when creating and maintaining complex codebases. TypeScript compiles to JavaScript and so can run in any modern web browser.

#### 3.2.2 User Interface

The Webina web app leverages Vue.js 2.6.10 (https://vuejs.org/), an open-source JavaScript framework for building single-page web applications with consistent user interfaces. Vue.js allows web programmers to create their own reusable HTML-like components (e.g., text fields, buttons, etc.). The properties and values of each component may depend on user-initiated actions (e.g., a “Submit” button may only be enabled once the user has entered text into an associated form). With each user action, Vue.js automatically updates all dependent components throughout the entire app. The Vue.js framework thus greatly simplifies the process of creating complex web apps with many interconnected features.

We also use the open-source BootstrapVue library 2.0.2 (https://bootstrap-vue.js.org/) to ensure that all user-interface components have a consistent appearance. BootstrapVue provides a number of broadly useful Vue.js components (e.g., buttons, check boxes, pop-up message boxes, etc). These components are styled according to the Bootstrap4 framework (https://getbootstrap.com/), which provides professionally designed templates that dictate components’ color, size, and typography.

We also created a custom molecular-visualization Vue.js component that leverages the 3Dmol.js JavaScript library [8]. This component allows the Webina web app to display macromolecular and small-molecule structures in the browser, as required for setting up Webina runs and analyzing Webina output. 3Dmol.js uses native web technologies to display these structures without requiring users to install any program or browser plugin.

#### 3.2.3 Build Process

We use Webpack, an open-source module bundler, to organize and assemble all custom and third-party libraries and files (https://webpack.js.org/). Webpack automatically builds the Webina web app by copying required files to their appropriate locations, combining files where possible, removing unnecessary/unused code, etc. Applying Google’s Closure Compiler is a critical component of this build process (https://developers.google.com/closure/compiler). The Closure Compiler automatically analyzes and rewrites our TypeScript/JavaScript code to reduce file sizes and speed execution.

### 3.3 Preparing a Benchmark Set

#### 3.3.1 Preparing Protein-Receptor Files

To compare Webina and Vina, we created a benchmark set of five protein/ligand complexes. To prepare the protein receptors for docking, we first removed all water molecules and ions. We then submitted the protein structures to the PDB2PQR server [9], a free resource that adds hydrogen atoms per a user-defined pH while simultaneously optimizing the hydrogen-bond network. We used the default PDB2PQR settings, which assign protonation states at pH 7.

We next used Open Babel [10] to convert the output receptor PQR files back to PDB. Vina requires input receptor files to be in the PDBQT format, a PDB-like format that additionally includes atom types and partial atomic charges. We used MGLTools 1.5.6 (http://mgltools.scripps.edu/), a desktop program created by the same group that created Vina, to convert the PDB files to PDBQT. We note that Open Babel can also convert files to the PDBQT format, but it does not always assign the same partial atomic charges and ligand rotatable bonds that MGLTools does.

One of the benchmark protein receptors, COX-2, contains heme groups. We similarly used MGLTools 1.5.6 to convert PDB files containing these heme groups to the PDBQT format. We appended these PDBQT-formatted files to the COX-2 PDBQT file prior to docking.

#### 3.3.2 Preparing Ligand Files

We used the DrugBank database [11] to obtain the SMILES strings of each input ligand. We then used the open-source program Gypsum-DL [12] to generate protonated ligand models from these SMILES. For each input ligand, Gypsum-DL can generate multiple output PDB files that collectively account for alternate ionization, tautomeric, chiral, cis/trans isomeric, and ring-conformational states. To simplify our analysis, we instructed Gypsum-DL to generate only one molecular variant per input molecule, protonated as appropriate for pH 7. We again converted the Gypsum-DL output PDB files to the PDBQT format using MGLTools 1.5.6.

#### 3.3.3 Webina and Vina Docking

We docked each ligand into its respective receptor using both Webina and Vina 1.1.2. In each case, the docking box was centered on the geometric center of the corresponding crystallographic ligand, calculated using the *scoria* Python library [13]. All docking boxes were 20 Å × 20 Å × 20 Å, cubed. Each docking run used four CPUs, an exhaustiveness setting of eight, and a consistent random seed of 123456789. We measured the root-mean-square deviation (RMSD) between each top-scoring docked pose and the corresponding crystallographic ligand using *obrms*, a program included in the Open Babel package [10].

### 3.4 LARP1 and PARP-1 Docking

We used a similar protocol to prepare La-related protein 1 (LARP1) and poly [ADP-ribose] polymerase 1 (PARP-1) receptors and ligands for Webina docking. For LARP1 docking, we used the 5V87:B structure, which captures LARP1 DM15 bound to m^7^GpppC [14]. The m^7^GpppC and m^7^GpppG ligands were prepared using Gypsum-DL as above, except the phosphate-bound hydrogen atoms were manually removed. The ligands were docked into a docking box with dimensions 16 Å × 17 Å × 17 Å, positioned so as to encompass both the 7-methylguanosine (m^7^G)- and +1C-binding pockets. To accommodate the flexibility of the triphosphate moiety, we ran Webina docking using an exhaustiveness setting of 20. Where two poses had the same predicted docking score, we favored the pose that placed the m^7^G cap in the cap-binding pocket.

For PARP-1 docking, we docked the known inhibitor pamiparib, currently in clinical trials [15], into three *holo* PARP-1 structures: 4HHY:A [16], 4R6E:A [17], and 4RV6:A [17]. As positive controls, we also docked the cognate ligands of these structures: a benzo[*de*][1,7]naphthyridin-7(8*H*)-one derivative, niraparib, and rucaparib, respectively. All PARP-1 Webina runs used an exhaustiveness setting of 8. Docking boxes were centered on the PARP-1 catalytic domain. We used boxes with dimensions 12 Å × 20 Å × 14 Å, 15 Å × 20 Å × 14 Å, and 16 Å × 20 Å × 14 Å for the 4HHY:A, 4R6E:A, and 4RV6:A structures, respectively.

## 4 Results and Discussions

### 4.1 Webina Library

Webina is a library for performing ligand docking in the web browser. It is built on the AutoDock Vina 1.1.2 codebase, compiled to WebAssembly. Web-app developers can incorporate the library into their own CADD apps. The Webina git repository includes a simple example HTML file that demonstrates use.

Vina command-line parameters are accessible via the Webina library, with the exception of the parameters used to specify input files. For security reasons, web browsers do not allow JavaScript to directly access users’ file systems. Instead, the contents of local files must be read through a file input element, using a JavaScript FileReader object. To overcome this limitation, the Webina library accepts text strings containing the contents of the input PDBQT files, rather than the file paths normally specified via Vina parameters such as --receptor and --ligand.

### 4.2 Webina Web App

#### 4.2.1 Input Parameters Tab

To accommodate users who are not web developers, we have incorporated the Webina library into a Webina web app that includes additional tools for setting up Webina runs and visualizing docking results. On first visiting the Webina web app, the user encounters the “Input Parameters” tab. This tab includes several subsections that are useful for setting up a Webina run (Figure 1).

**Figure 1:**
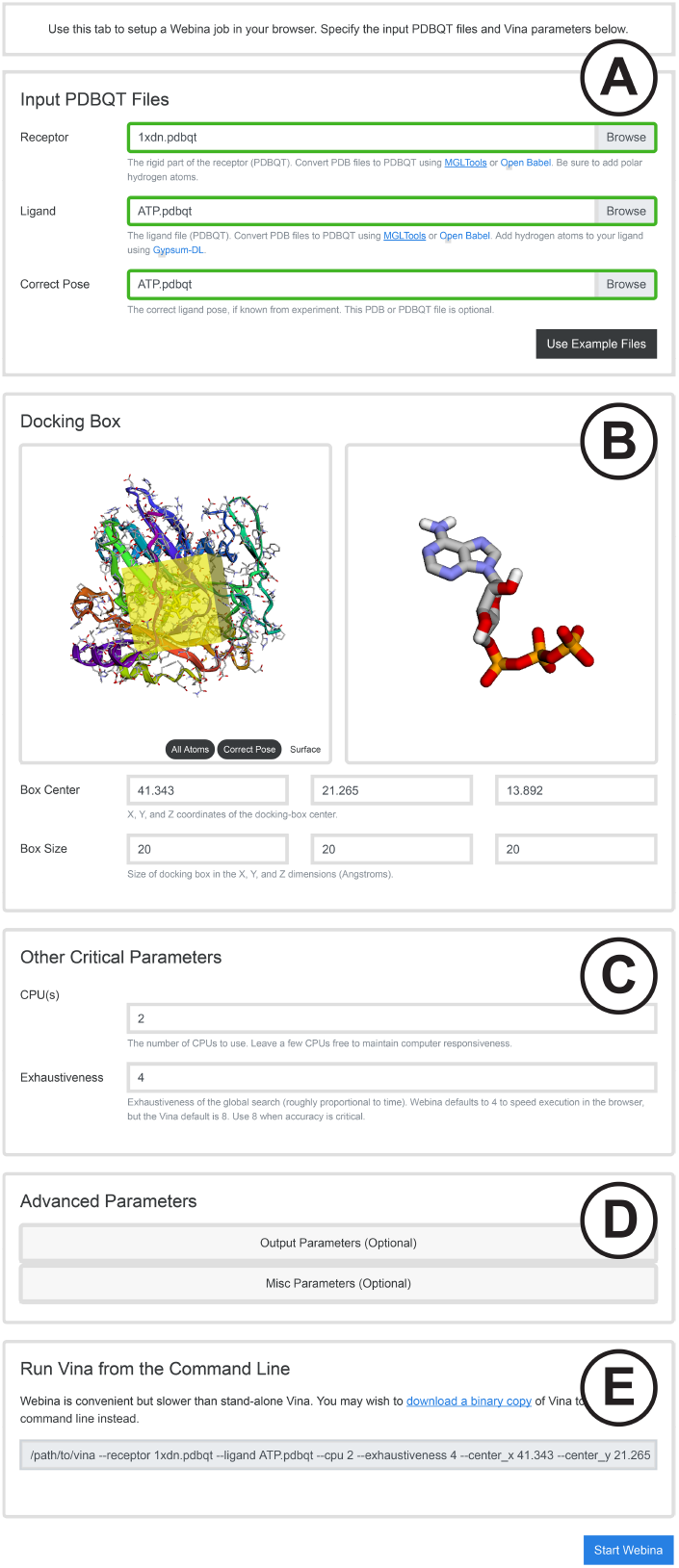
The Input Parameters tab includes the A) Input PDBQT Files, B) Docking Box, C) Other Critical Parameters, D) Advanced Parameters, and E) Run Vina from Command Line subsections.

##### Input PDBQT Files

The “Input PDBQT Files” subsection allows the user to select their receptor and ligand files (Figure 1A). As is the case with command-line Vina, these files must be in the PDBQT format. The user can also optionally specify a known-pose PDB or PDBQT ligand file. This file includes the ligand in its experimentally determined, correct bound pose (e.g., per X-ray crystallography or NMR). The known-pose file plays no role in the docking calculation; rather, it serves as a positive-control reference for evaluating Webina-predicted ligand poses. In our experience, it is often helpful to first benchmark Webina (or Vina) against a known ligand before using the program to evaluate compounds with unknown poses and binding affinities.

Some users may wish to test Webina without having to provide their own input files. The optional “Use Example Files” button automatically loads in example receptor, ligand, and known-pose files.

##### Docking Box

The “Docking Box” subsection allows users to specify the region of the receptor where Webina should attempt to pose the ligand (Figure 1B). This box-shaped volume is typically centered on a known pocket or cavity where a small-molecule ligand might reasonably bind.

To simplify the process of selecting a docking box, the Webina web app automatically displays 3D models of the user-specified receptor and ligand using the 3Dmol.js JavaScript library [8]. By default, the receptor and ligand are displayed using cartoon and sticks representations, respectively. The user can toggle a surface representation as required to identify candidate receptor pockets. A transparent yellow box is superimposed on the structures to indicate the docking-box region.

When the user clicks the atoms of the receptor model, the Webina web app recenters the docking box on the selected atom. Users can also adjust the location and dimensions of the box using text fields below the molecular visualization.

##### Other Critical Parameters

The “Other Critical Parameters” subsection allows the user to specify the number of CPUs and the exhaustiveness setting (Figure 1C). We chose to set these two parameters apart because they are particularly important in a browser-based setting. Users expect command-line tools to consume substantial computer resources, but they do not expect web apps to do so. By default, Vina uses all available CPUs and an exhaustiveness setting of eight. Webina has the same ability to consume CPUs and memory, but many users will wish to adjust these parameters to avoid impacting the performance of other programs and browser tabs.

##### Advanced Parameters

The “Advanced Parameters” subsection allows users to specify the many additional parameters that are also available via command-line Vina (Figure 1D). In our experience, it is rarely necessary to adjust these parameters, so they are hidden by default.

##### Run Vina from Command Line

The “Run Vina from Command Line” subsection aims to help Vina users who wish to use the Webina web app to setup their docking boxes and user parameters (Figure 1E). A text field provides a mock example of how to use command-line Vina with the specified parameters. Users can copy this example, modify it as needed, and paste it into their command-line terminals to run the desired calculation with the standard Vina executable.

##### Starting the Webina Calculation

Once users click the “Start Webina” button, the Webina app will switch to the “Running Webina” tab while Webina executes. During this process, the user-specified receptor/ligand files and parameters are never transmitted to any third-party server. The calculations run entirely in the user’s browser itself. The Webina browser tab may become unresponsive during execution as a result. Once the calculation is complete, the Webina web app will switch to the “Output” tab (described below) where users can visualize the docking results.

#### 4.2.2 Existing Vina Output Tab

Beyond serving as a platform for running the Webina library, our Webina web app provides a useful interface for visualizing docking calculations. The “Existing Vina Output” tab allows users to load and visualize the results of previous Webina and Vina runs, without having to rerun the calculations. Users must specify the existing receptor and Webina/Vina output file they wish to visualize. They can also optionally specify a known-pose ligand file for reference. Users who wish to test the web app without providing their own files can click the “Use Example Files” button. Otherwise the “Load Files” button will open and visualize the specified files.

#### 4.2.3 Output Tab

The “Output” tab allows users to visualize their Webina docking results (Figure 2). The same tab also displays the output of any previous Webina/Vina calculations that the user specifies via the “Existing Vina Output” tab.

**Figure 2:**
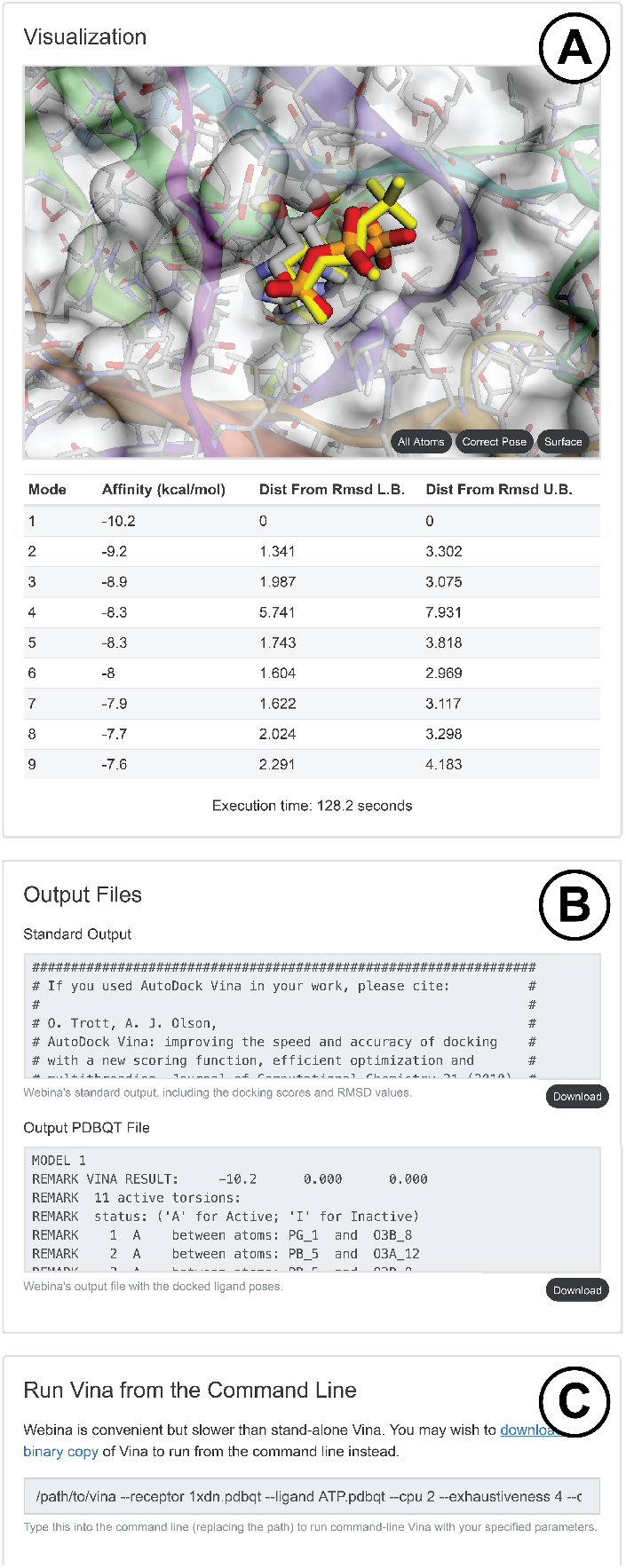
The Output tab includes the A) Visualization, B) Output Files, and C) Run Vina from Command Line subsections.

##### Visualization

The “Visualization” subsection uses 3Dmol.js to display the receptor and docked molecule in cartoon/surface and sticks representation, respectively (Figure 2A). If the user has specified a known-pose ligand file, that pose is also displayed in yellow sticks. Like Vina, Webina predicts several poses per input ligand. A table below the visualization viewport lists each pose together with associated information such as the docking score. Clicking on a table row updates the 3D view with the specified pose so users can easily examine all predicted poses.

##### Output Files

The “Output Files” subsection shows the text contents of the Webina output files (Figure 2B). An associated “Download” button allows users to easily save those files.

##### Run Vina from Command Line

Similar to the “Input Parameters” tab, the “Output” Tab also includes a “Run Vina from Command Line” Subsection (Figure 2C). This section makes it easy for users to reproduce Webina’s results using stand-alone Vina. It also reminds users what parameters they selected to generate the displayed Webina output.

#### 4.2.4 Start Over Tab

The “Start Over” tab displays a simple button that allows the user to restart the Webina app. A warning message reminds the user that they will loose the results of the current Webina run unless they have saved their output files.

### 4.3 Benchmarks

To compare Webina and Vina output and run times, we created a benchmark set of five protein/ligand complexes. We intentionally selected complexes whose ligands are clinically approved and whose proteins have diverse structures and functions: HMG-CoA reductase/rosuvastatin (PDB ID: 1HWL [18]), factor Xa/apixaban (PDB ID: 2P16 [19]), COX-2/celecoxib (PDB ID: 3LN1 [20]), HIV protease/darunavir (PDB ID: 4LL3 [21]), and cereblon/S-lenalidomide (PDB ID: 4TZ4) (Table 2).

**Table 2:**
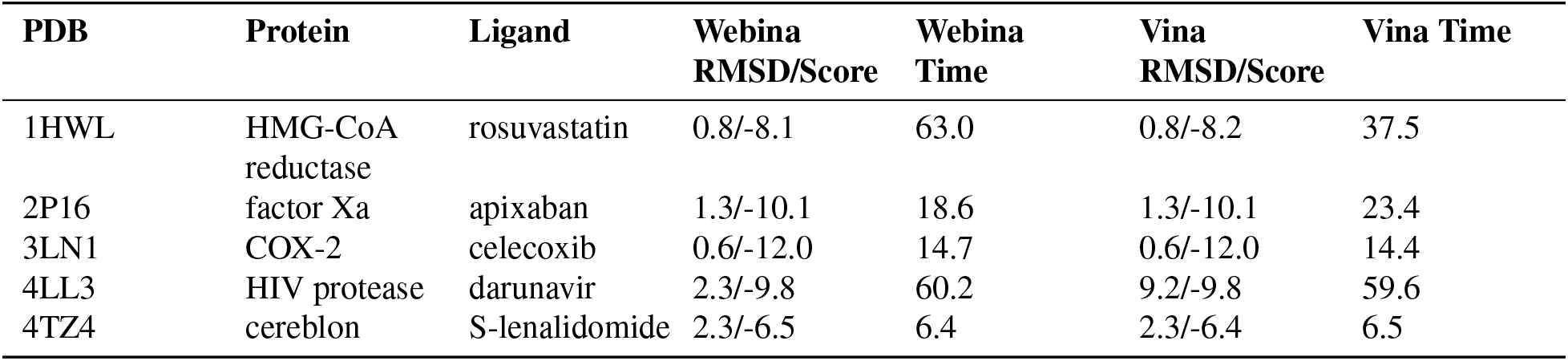
Webina/Vina Benchmarks. For both Webina and Vina, we considered only the top-scoring pose. RMSD refers to the root-mean deviation from the crytstallographic pose (in Å), calculated using the Open Babel *obrms* program. Scores are reported in kcal/mol. Times are reported in seconds.

#### 4.3.1 Poses and Scores

Webina and Vina 1.1.2 produced comparable ligand poses and docking scores, as expected given that both are compiled from the same codebase (Table 2). Given that we used the same random seed for both Vina and Webina, we were surprised that the output wasn’t identical. We believe that rounding differences are responsible for the minor discrepancies. Per the official WebAssembly Specification Release 1.0 (Draft, Oct 29, 2019), Wasm deviates slightly from the IEEE 754-2019 standard for floating-point arithmetic.

Despite these minor differences, Webina and Vina produced similar docking scores. For three of the five test complexes, the scores were identical. For the remaining two, the scores differed by only 0.1 kcal/mol (Table 2). Webina and Vina also produced similar docked poses. In four of the five test cases, the top Webina and Vina poses differed from each other by at most 0.22 Å.

The HIV protease/darunavir complex (PDB ID: 4LL3 [21]) was a curious exception. The 4LL3 crystal structure describes two distinct protein/ligand conformations that are both presumably valid. These two conformations capture two very different ligand poses, with accompanying rotomeric differences in the ligand-interacting amino acids. We used the first protein conformation for all test runs and discarded the second. The top Vina pose (−9.8 kcal/mol) nevertheless most closely matched the ligand from the second conformation. In contrast, the second Vina pose (−9.7 kcal/mol) matched the correct conformation (2.3 Å). Because of the rounding differences described above, Webina instead assigned both these poses the same score (−9.8 kcal/mol) and happened to list the more correct pose first.

We expect that the choice of random seed will have a far greater impact on the docking results than any slight differences in rounding. For example, when we reran the HIV protease/darunavir docking using a random seed of 1000, Vina reversed the ranking of the first and second pose, now placing the more correct pose in first place.

#### 4.3.2 Execution Speed

Webina took slightly longer to finish than did Vina 1.1.2, as expected given that it is a Wasm-compiled program. We tested Webina and Vina 1.1.2 on a MacBook Pro (15-inch, 2018) running macOS Mojave 10.14.5 (2.9 GHz Intel Core i9 processor and 32 GB 2400 MHz DDR4 memory). On average, Webina took roughly 10% longer to dock the compounds than did Vina. When running a large-scale virtual screen, we recommend using the faster and more scalable command-line version of Vina. But when docking only a few compounds, Webina is an ideal, user friendly solution that requires no installation or command-line use.

### 4.4 Examples of Use

#### 4.4.1 LARP1 Specificity

To demonstrate utility, we used Webina to study ligand binding to LARP1, a potential anti-cancer drug target [22–24]. LARP1 regulates the translation of key proteins required for ribosome biogenesis [25–29]. The mRNAs that encode these proteins have a characteristic 5’ terminal oligopyrimidine (TOP) motif, i.e., a cytosine invariably occupies the +1 position adjacent to the m^7^G cap, followed by a tract of pyrimidines [30]. TOP mRNAs thus differ from most protein transcripts, which typically have a purine in the +1 position. The LARP1 DM15 region binds both the m^7^G cap and the invariant +1 cytosine (+1C) characteristic of TOP motifs [14, 31]. We have previously shown using mutagenesis that the +1C pocket drives LARP1-TOP specificity [32].

We used Webina to see whether we could predict LARP1 TOP-motif specificity *in silico*. We docked both the m^7^GpppC and m^7^GpppG dinucleotides into the 5V87:B DM15 structure, which was co-crystallized with bound m^7^GpppC [14]. In harmony with previous experimental studies [14, 31], Webina scored the top m^7^GpppC pose slightly higher than the top m^7^GpppG pose (−6.2 and −5.9 kcal/mol, respectively). Notably, the best m^7^GpppC pose captures many of the same interactions seen crystallographically (Figure 3A) [14]. The m^7^G cap forms hydrogen bonds with E886, as well as π-π and cation-π interactions with Y922 and Y883. Similarly, the +1C forms hydrogen bonds with R847 and π-π stacking interactions with Y883 and F844. In contrast, the best m^7^GpppG docked pose did not orient the +1G nucleotide such that its Watson-Crick face could interact with the +1C-pocket protein residues (Figure 3B).

**Figure 3:**
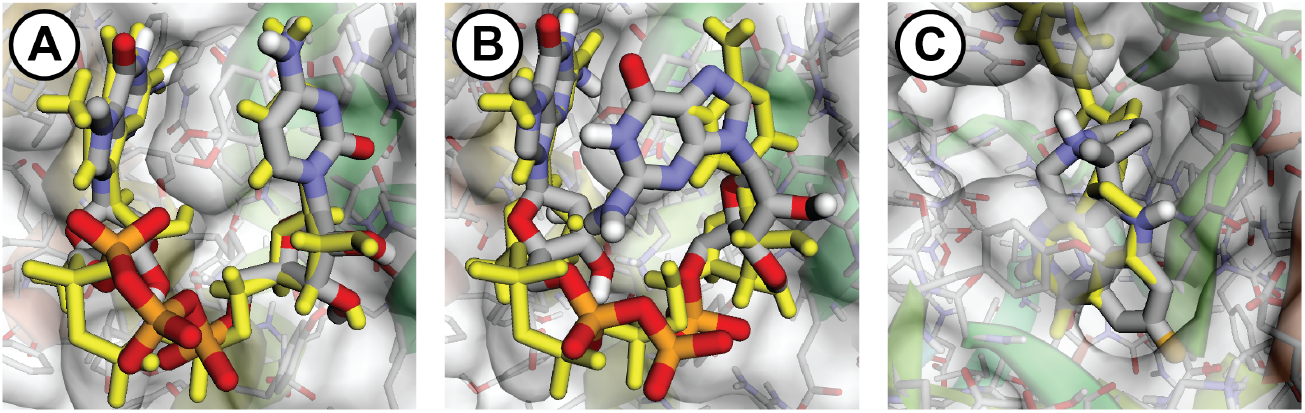
Screenshots of Webina docking results. Docked poses are shown in sticks representation, colored by element. Crystallographic poses are shown in yellow sticks. A) The m^7^GpppC/LARP1-DM15 docked pose recapitulates many of the hydrogen-bond, π-π-stacking, and cation-π interactions characteristic of the crystallographic pose. B) The m^7^GpppG/LARP1-DM15 docked pose did not orient the +1G Watson-Crick face towards the protein. C) The pamiparib/PARP-1 docked pose is similar to the crystallographic pose of a benzonaphthyridinone ligand.

We do not wish to overstate the utility of docking in general or Webina in particular. Docking certainly has its inaccuracies. For example, when we similarly docked the m^7^GpppC and m^7^GpppG dinucleotides into a LARP1 DM15 mutant engineered to specifically bind +1G transcripts (6PW3:C [32]), the poses were not nearly as good (data not shown). But, typical docking inaccuracies aside, Webina is still a powerful, user friendly tool for generating biologically relevant hypotheses.

#### 4.4.2 PARP-1 Ligand-Pose Prediction

As a second demonstration of utility, we used Webina to study PARP-1, a multifunctional nuclear protein involved in the base and nucleotide excision DNA repair pathways (BER and NER) [33]. PARP-1 is overexpressed in various carcinomas and is associated with more aggressive cancer and lower survival rates [34, 35]. BRCA-deficient carcinomas are especially susceptible to PARP-1 inhibition because they depend solely on BER for survival [36].

To verify that Webina can effectively identify PARP-1 ligands, we first used it to recapitulate the binding poses of three crystallographically characterized PARP-1 inhibitors: a benzo[*de*][1,7]naphthyridin-7(8*H*)-one derivative (PDB ID: 4HHY:A [16]), niraparib (PDB ID: 4R6E:A [17]), and rucaparib (PDB ID: 4RV6:A [17]). We docked each ligand into all three crystal structures. As expected, the best Webina-predicted binding affinity of each ligand was obtained when each was docked into its corresponding crystallographic conformation: the benzonaphthyridinone derivative, −14.7 kcal/mol; niraparib, −10.1 kcal/mol; and rucaparib, −10.8 kcal/mol. The heavy-atom RMSDs between the crystallographic and top-scoring docked poses were: the benzonaphthyridinone derivative, 2.35 Å; niraparib, 1.15 Å; and rucaparib, 0.77 Å.

Having verified that Webina is effective in this context, we next used it to predict the binding pose of pamiparib, a PARP-1/2 inhibitor currently in phase II clinical trials [15]. We docked pamiparib into the same three PARP-1 conformations. The PARP-1 structure associated with the top-scoring pamiparib pose (−11.3 kcal/mol) was 4HHY:A (Figure 3C) [16]. In this docked pose, pamiparib shares much in common with the crystallographic poses of the three positive-control ligands. All four overlay a benzamide substructure at the same location (Figure 3C), enabling hydrogen bonds with S904 and G863 and a π-π stacking interaction with Y907. To the best of our knowledge, there is no publicly available PARP-1/pamiparib crystal structure. This work thus demonstrates how Webina can be used to prospectively predict ligand poses, a critical component of many CADD workflows.

### 4.5 Comparison with Other Approaches

#### 4.5.1 Standard Vina Executable

Webina has several advantages over the standard Vina executable. Though code compiled to Wasm tends to run somewhat slower than natively compiled code, it runs entirely in the browser. And given that modern browsers already run on all major operating systems, Wasm-compiled programs are cross-platform by default. The Webina web app also provides a helpful GUI that command-line Vina lacks, making computer docking much easier for novices and students.

#### 4.5.2 Third-Party Graphical User Interfaces

Several groups have created GUI-based desktop applications for setting up, performing, and analyzing Vina results. Many of these interact with command-line Vina “under the hood.” For example, PyRx [3], now a commercial program, provides a helpful wizard-like interface for Vina docking. Programs such as DockingApp [4] and AUDocker LE [5] are somewhat similar (albeit less feature rich) open-source alternatives. Other GUI programs aim to simplify specific steps in the standard docking protocol. For example, MGLTools (http://mgltools.scripps.edu/), produced by the same group that created Vina, allows users to prepare ligand and receptor PDBQT files and to select an appropriate docking box. Popular molecular-visualization programs such as Chimera [37] and PyMOL [38] also include plugins (both commercial and free) for setting up and visualizing Vina docking runs [6, 7].

These tools are certainly useful, but they still require users to download and install separate executable files. Furthermore, some of these tools are costly, work on only some operating systems, and still suffer from usability despite their GUI implementations. In contrast, Webina works entirely in the browser on all operating systems and is as easy to use as visiting a standard web page.

#### 4.5.3 Server Applications

Recognizing the advantages of web-based docking, others have created server-based web apps that allow users to perform Vina docking on remote computer resources. MTiOpenScreen [39] is a good example of such an app. Though effective, the server-based approach requires app creators to maintain and secure the substantial remote resources needed to perform docking runs. Many users, especially those working in industry, may also be reluctant to upload their proprietary receptor and small-molecule structures to any third-party server. In contrast, Webina requires only one-way communication from the server to the browser. The browser downloads the Webina library just as it would an image or video. Webina then performs all docking calculations in the browser itself, without needing to transmit information about the docking input or output back to the server.

### 4.6 Conclusions

Computer docking is a common CADD technique. AutoDock Vina is a particularly popular docking program, but it requires download, installation, and command-line use. While CADD experts may well prefer command-line Vina, it is not ideal for those who wish to only occasionally dock compounds, or for students who do not yet have the required computer expertise.

Webina aims to address these challenges. The Webina library is built on the Vina 1.1.2 codebase, so the two programs produce comparable results. But Webina runs entirely within the web browser. Our associated Webina web app, which leverages the Webina library, also provides user-friendly tools for setting up docking calculations (e.g., identifying an appropriate docking box) and analyzing docking output (e.g., examining predicted binding poses). Aside from allowing users to dock compounds in their browsers, Webina is also a useful tool for examining the output of previously executed docking runs produced by Webina itself or command-line Vina.

Webina will be a useful tool for the CADD community. We release it under the terms of the Apache License, Version 2.0. A copy can be obtained free of charge from http://durrantlab.com/webina-download, and the web app is accessible at http://durrantlab.com/webina.

## 6 Acknowledgements

We acknowledge the University of Pittsburgh’s Center for Research Computing for providing valuable computer resources.

